# Novel Protein Structure Validation using PDBMine and Data Analytics Approaches

**DOI:** 10.1101/2025.02.07.637116

**Authors:** Niharika Pandala, Katherine G. Brown, Homayoun Valafar

## Abstract

Protein structure prediction is essential for understanding biological functions and advancing drug development. Although experimental techniques like NMR, X-ray crystallography, and cryo-EM provide valuable insights, they are expensive and time-consuming, prompting reliance on computational approaches. AlphaFold2 revolutionized protein model predictions accuracy in 2020. However, limitations remain in the prediction of novel proteins, complex conformations, and mutations. To address these challenges, we leverage PDBMine software and machine learning for advanced data analytics. This approach detects and corrects structural inaccuracies, calculates fitness scores, and enhances model reliability, accelerating drug discovery and therapeutic breakthroughs by bridging gaps in current protein prediction capabilities.

## 1 Introduction

A protein’s significance in biological processes requires understanding the link between its three-dimensional structure and biological function (Taufer et al., 2005). Understanding protein structure is crucial for altering or regulating protein function in disease treatment and drug development. However, the conventional experimental methods determining protein structures via Nuclear Magnetic Resonance (NMR) spectroscopy, X-ray crystallography, and cryo-electron Microscope(cryo-EM) pose considerable difficulties due to their time-consuming nature, high costs, and need for specialized expertise.

To address these challenges, the scientific community is increasingly exploring computational techniques as a substitute for determining protein structures. These methods use the Protein Data Bank (PDB) (Berman et al., 2003) as a primary resource to bridge the gap between a protein’s sequence and structural configuration. Recent advances in artificial intelligence (AI) and machine learning (ML) have significantly improved protein structure prediction accuracy through tools such as ROSETTA3 (Leaver-Fay et al., 2011), I-TASSER (Yang J et al., 2016), AlphaFold2 (Jumper et al., 2021), and “Artificial Neural Network approaches to enhancing the correlation between protein energetics and backbone structures” (Fawcett, Timothy M et al., 2013). These algorithms use approaches rooted in similarity detection like homology, de novo, and ML modeling, thereby increasing structure prediction accuracy (Yang Z. et al., 2023).

A notable breakthrough was observed in this domain in 2020 with the success of AlphaFold2 (Jumper et al., 2021) in the biennial Critical Assessment of Structure Prediction (CASP) competitions (Kryshtafovych A. et al., 2021) that facilitates advancement in computational protein structure prediction. AlphaFold’s neural network-based models provide a unique technique for predicting protein structures, establishing a new industry benchmark for protein modeling accuracy and outperforming other existing methods. Despite its remarkable advancement, AlphaFold2 still faces limitations, particularly in determining complex proteins, novel proteins, and loop structure (Stevens et al., 2022), only predicting a single protein conformation, and failing to predict the defect caused by mutations (Buel et al., 2022). Our proposed methodology presents an advanced data analytics approach that is pivotal in detecting and correcting structural discrepancies in computationally predicted protein models by leveraging machine learning and integrating statistical data from the PDB via PDBMine (C. Cole et al., 2019). This comprehensive approach not only increases model precision but also offers correction that leads to deeper insights into molecular interactions, instilling confidence in its potential for novel drug discoveries and innovative treatments for diseases.

## 2 Background

PDB is a crucial resource for recording 3D structural information about biological molecules. About 187,089 proteins are in the PDB (Berman H.M. et al., 2007). PDB usage has risen significantly because computational techniques facilitate structure modeling and prediction using template-based modeling (TBM). Such models accuracy heavily depends upon the relationship between the protein sequences of unknown proteins and those of known structures in PDB, making homology searches and comparative modeling easier (Pearce R et al. 2021). The predictive capacity accelerates structural biology discoveries and is essential for comprehending the role of unknown or novel proteins. Computing techniques have aided in predicting the structure of related proteins, further enhancing our knowledge of protein families, evolutionary links, and comparative modeling.

Overall, predicting possible protein structure models using computational approaches has simplified a variety of scientific endeavors, from experimental design optimization to making research on novel proteins more affordable and accessible to a broader scientific community; this has allowed for time and resource savings by concentrating efforts on the most promising direction. Researchers can swiftly process and evaluate large amounts of data using computational methods for a high-throughput study of protein structures. Large-scale initiatives like investigating entire proteomes or performing genome-wide association studies (GWAS) depend heavily on this efficiency. Hence, by developing these predictive technologies, we address the disparity between the number of known protein sequences and experimentally predicted structures (Pearce et al. 2021) to better understand the protein structure and function paradigm.

## 3 Methodology

The proteins utilized in this investigation were sourced from the Critical Assessment of Structure Prediction (CASP14) dataset, obtained from the official CASP prediction center website. CASP14 encompasses an automatic evaluation center housing all proteins submitted as target lists alongside predictions from 113 participating groups during the 2020 CASP cycle. Following the competition, each target protein’s finalized experimental structure was deposited into the Protein Data Bank (PDB) for subsequent analysis, serving as the ground truth for comparison against computational predictions. For this study, we worked with ten protein models; specifically, we selected five well-predicted models, and five poorly predicted models based on AlphaFold2 prediction. Since AlphaFold2 consistently outperformed other CASP competitors for most models, exhibiting greater structure prediction accuracy. However, by evaluating its successes and failures, we hope to better understand the factors impacting AlphaFold2 when its forecasts were inaccurate. This comparison analysis offers insightful information about areas where protein structure prediction methodologies can be improved.

### 3.1 Introduction to PDBMine Analysis

The initial step in utilizing the PDB is to extract the atomic coordinates from the PDB files to glean insights into the structural characteristics of CASP proteins. To reduce the dimensionality of the data by an order of magnitude set for each PDB file, PDBMine software was used to derive all possible torsion angle subspaces of the given protein, while Python and Biopython facilitated the determination of dihedral angles constituting the protein backbone.

PDBMine software enabled efficient deduction of torsion angles by querying the PDB database with a specified protein sequence and user-defined k-mer size. This approach facilitated the retrieval of torsion angles associated with each amino acid in the given sequence motifs across the database, thus enabling a more intricate structural analysis. Notably, the choice of k-mer size specified by the user significantly influences the results specificity and relevance; larger k-mers offer greater precision while yielding fewer results, whereas smaller k-mers provide broader insights with potentially less direct significance. We used kmer sizes of 4mers, 5mers, and 7mers in our analysis. Subsequently, the data from each protein run encompassed all possible torsion angle subspaces for individual amino acids within the protein sequence for a given kmer, allowing comprehensive exploration of structural configurations.

Following data extraction, a filter was implemented to have torsion angles only in the Ramachandran subspace with a range of -180 to 180 degrees. The subsequent analysis involved plotting the profile of all possible phi and psi configurations for each amino acid residue, yielding Ramachandran plots. These plots allowed us to discern clusters of preferred torsional configurations across extensive datasets, offering valuable insights into the structural flexibility and constraints of the proteins. Most of the computational analyses were conducted utilizing Python and Biopython, ensuring the robustness and reproducibility of our results.

### 3.2 Ramachandran Space and Likelihood Analysis using KDE

This proposed methodology harnesses computational strategies to investigate clustering phenomena within the Ramachandran plot, a crucial protein structure prediction and refinement tool. The Ramachandran plot provides a foundational map for analyzing protein structure by illustrating sterically allowed and disallowed regions based on the phi (φ) and psi (ψ) torsion angles. These regions, which include energetically favored conformations such as alpha helices and beta sheets, are essential for understanding protein folding dynamics and structural propensities (Ramachandran et al., 1963). Steric constraints prevent certain torsion angles due to atomic clashes, and residues falling into disallowed regions may indicate errors in the protein model. By incorporating experimentally determined torsion angles as benchmarks, this approach allows for a robust comparison of computational models against actual protein structures, ensuring model accuracy and structural integrity.

We employed Kernel Density Estimation (KDE) to refine structural predictions further using Silverman’s method (B.W. Silverman, 1986). This technique estimates the distribution of structural features such as torsion angles or distances between atoms by smoothing the observed data from PDBMine, particularly from k-mer sizes of 4mers, 5mers, and 7mers. By providing an optimal bandwidth for KDE, Silverman’s method strikes a balance between detail and noise in the density estimates, thereby enabling a clearer visualization of preferred protein conformations. This approach enhances the accuracy of structural predictions by capturing underlying trends in the protein’s geometry and enables the identification of statistically significant conformations. This, in turn, facilitates the identification of structural preferences, guiding more informed decisions in protein modeling and drug design.

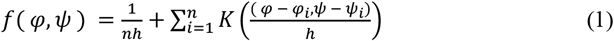

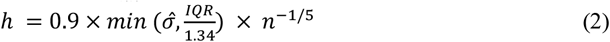

KDE is determined using equation 1, where *f*(*φ, ψ*) the estimated density at a specific pair of torsion angles ϕ and ψ. Here, *n* is the number of data points (torsion angles) for the protein, and *h* is the bandwidth that controls the smoothness of the density estimate. *K* is the kernel function, typically Gaussian, that smooths the data over the angle space. The Silverman method is used in equation 2 to select the optimal bandwidth *h* for KDE, where 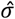 is the standard deviation of the torsion angles, *IQR* is the interquartile range of the torsion angles and *n* is the number of torsion angles (data points).

### 3.3 Considering Other Potential Clusters Using Threshold Analysis

In contrast to the approach outlined in section 3.3, which focuses on one ideal cluster of protein data points, this method takes a more flexible approach. It leverages the same KDE profiles but identifies additional clusters with plausible torsion angles for each residue, allowing for more adaptability in the fitness scores. This is especially helpful when a torsion angle point is equally likely to appear in more than one cluster, as experimental data often shows multiple clusters. Instead of penalizing predictions for not falling into the most probable cluster, this approach accepts alternative clusters when justified.

To ensure only high-likelihood clusters are considered, a threshold is established. We aim to accept only clusters with significant likelihood rather than accepting any point across the entire Ramachandran plot. After plotting the Ramachandran subspace for a given residue, a 3D KDE is generated, where the third dimension represents the likelihood of torsion angles. The threshold is set at 0.004% of the likelihood of the most probable torsion angle. This value was derived from studying experimental X-ray data for the proteins in the study. With this threshold, the Ramachandran space is narrowed down to regions with higher likelihood probabilities, accounting for the presence of multiple plausible clusters.

Once the 0.004% threshold is applied, a binary mask is created for the entire interpolated KDE Ramachandran space with a resolution of 500×500. The algorithm checks whether the likelihood falls below or above the threshold for each point in this grid. Points below the threshold are assigned a value of 0, while points above are assigned 1, creating a binary mask that clearly identifies acceptance and rejection regions. After the mask is in place, we quantify the percentage of space accepted and rejected for each residue across the 10 CASP target protein models and their 113 predictions.

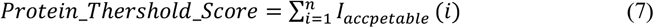

Where I_*acceptable*_(*i*) = 1 if residue *i* falls within the acceptable region and I_*acceptable*_(*i*) = 0 if residue *i* falls outside the acceptable region. This scoring method offers flexibility by considering possible clusters rather than strictly probabilities, with an average acceptance rate of 20% per residue. A protein’s score is calculated by summing the number of residues that fall within the acceptable region, as shown in equation 7. In the case of X-ray structures, the score will theoretically be equal to *n*, the total number of residues. A model is considered accurate if its score is closer to *n*, indicating better agreement with the experimentally validated structure.

### 3.4 Neighboring points Analysis

In certain cases, involving specific protein residues, the Ramachandran plot contains a limited number of data points, indicating sparse experimental evidence for particular regions within the Ramachandran subspace. An alternative approach was developed to assess the proximity of neighboring torsion angle points to address these cases. This section utilized a dataset comprising twenty-two domain proteins, with an equal split between well-predicted AlphaFold2 models and poorly predicted AlphaFold2 models. This balanced selection ensures a comprehensive analysis of AlphaFold’s performance across a spectrum of prediction quality.

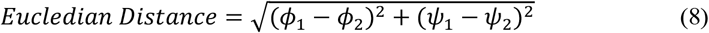

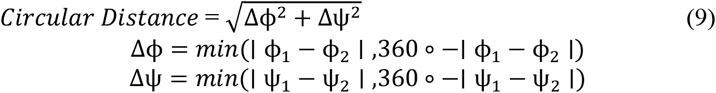

This method involved analyzing the torsion angle by defining a 5-degree radius around the point of interest in the Ramachandran space. Data from PDBMine, including 4, 5, and 7-mer window sizes, were utilized to calculate the Euclidean distance using equation 8. Then, circular distance was implemented due to the periodicity of the angular space in the Ramachandran plot using equation 9. These distances were calculated between the queried PDBMine point (ϕ_1,_ ψ_1_) from the Protein Data Bank (PDB) and the predicted point (ϕ_2_, ψ_2_) of the computationally modeled structure. Various circular cutoffs were implemented after determining the circular distance, ranging from 5 to 55 degrees in 5-degree increments, enabling a systematic evaluation of structural deviations.

Input features for each category representing the lack of nearby data points surrounding isolated predicted locations on the Ramachandran plot were used to train the decision tree classifier. For each prediction model, these features were obtained by counting the number of isolated spots with no adjacent data between radius increments ranging from 5 to 55 degrees (in 5-degree steps). We examined five prediction models produced by AlphaFold2 for every protein, which captured the spatial isolation of sites at different radial thresholds. The decision tree could categorize models as acceptable or unacceptable based on their RMSD with a specified RMSD of 6 by integrating this spectrum of isolation data, giving detailed insights into the relationship between the geographical distribution around isolated sites and structural accuracy. With Leave-One-Out Cross-Validation offering reliable confirmation of predicted performance, this method improves the classifier’s capacity to identify structural dependability across various protein prediction models.

## 4 Results and Analysis

### 4.1 Ramachandran space and Likelihood Analysis KDE

A detailed comparison between predicted and experimental torsion angles was performed using Ramachandran plots and 3D likelihood projections. Ramachandran plots analyzed residue by residue using PDBMine’s k-mer results, which reveal key scenarios. In the best-case scenario, the ground truth, PDBMine, and AlphaFold2 predictions align, showing a solid match between predicted and experimental structures. In these cases, the Ramachandran space is densely populated in regions corresponding to expected secondary structures, reinforcing the model’s accuracy.

Notably, using larger window sizes tends to provide more robust structural insights. For example, **Figure 1b** from Protein T1030-D2 demonstrates that the X-ray structure aligns with the cluster peak in the 5-mer pool, with high-intensity areas in the 3D-KDE map indicating a high likelihood of observing relevant torsion angle pairs in the alpha helix domain. This suggests that larger window sizes enhance prediction accuracy by better capturing the underlying structural patterns.

**Fig. 1.**
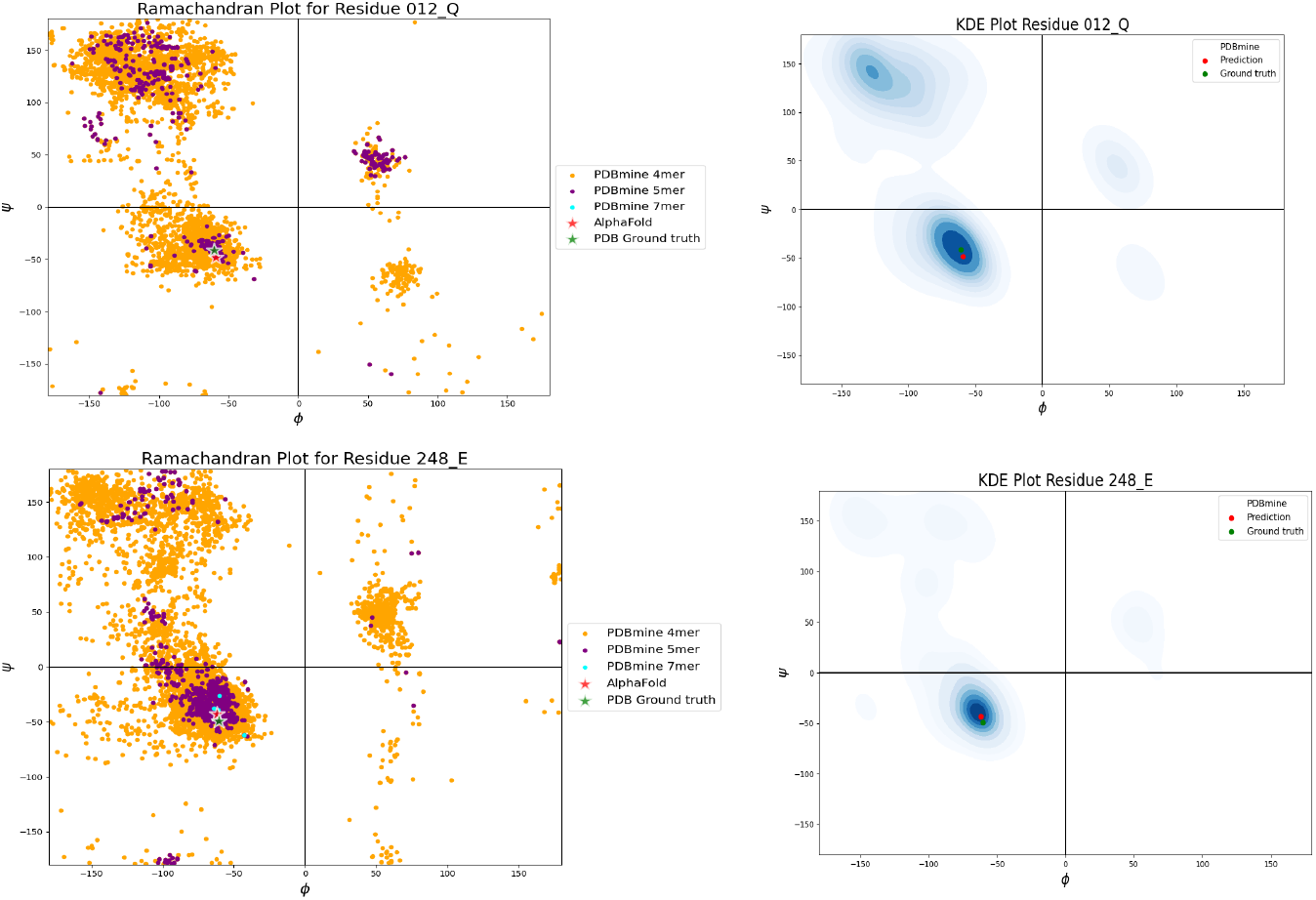
Ramachandran plots on the left depict PDBMine results for window sizes of 3mers (orange), 4mers (purple), and 7mers (cyan), with the X-ray torsion angle marked by a green star and the predicted AlphaFold2 model by a red star. The corresponding 3D projections on the right show the likelihood density, where PDBMine is represented as a density plot and experimental truth and prediction are marked by green and red points, respectively. 1a corresponds to T1024 residue 12 (Glutamine), and 1b to T1030-D2 residue 248 (Glutamic acid).

PDBMine provides a clearer understanding of model accuracy for specific residues such as T1024’s Glutamine and T1030-D2’s Glutamic acid. In the left panels of **Figure 1** to **Figure 3**, PDBMine results for window sizes of 3mers, 4mers, and 7mers, considering the neighboring residues based on the window size that populate the torsion angle space for each given residue of the sequence. Further, the green star also marks X-ray torsion angles, and AlphaFold2 prediction is marked by a red star. The right panels extend the analysis by incorporating a 3D likelihood map, where the experimental and predicted torsion angles are compared to generated density plots.

On the other hand, we observe instances where the experimental ground truth and PDBMine agree, but the predicted model, particularly AlphaFold, does not align. Since AlphaFold2 has generally outperformed most predictive models, the figures in this section focus on it as the primary predictive model. These represent cases where the discrepancies in specific residues, solvable through PDBMine, could improve the accuracy of the predicted protein structure. For example, in **Figure 2**, both T1032-D1 and T1038 show agreement with the PDBMine cluster, whereas AlphaFold2 selects a different cluster. By addressing these differences, PDBMine can aid in refining and enhancing the structure prediction of these models.

**Fig. 2.**
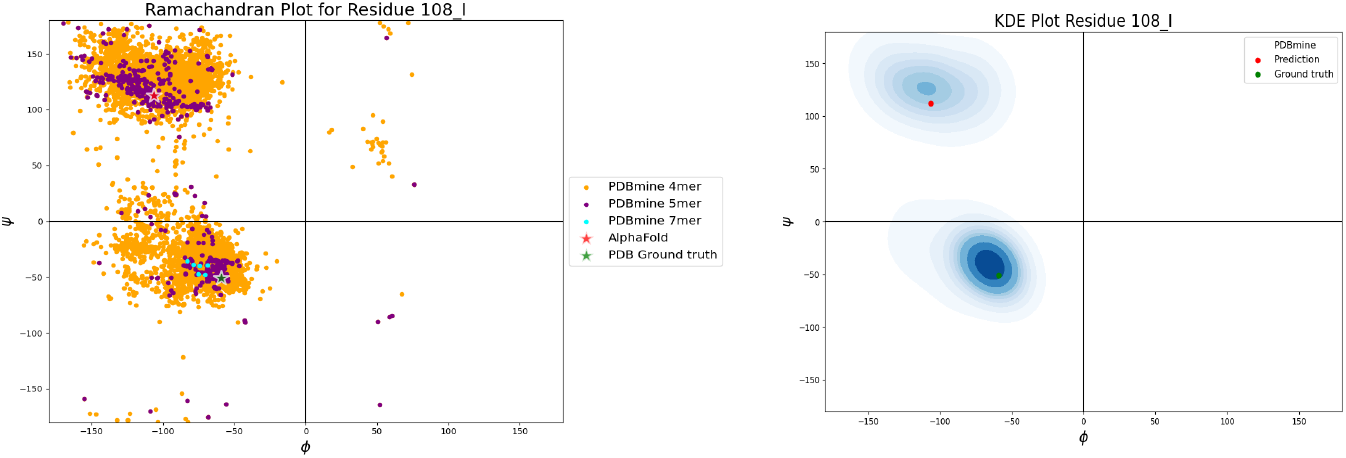
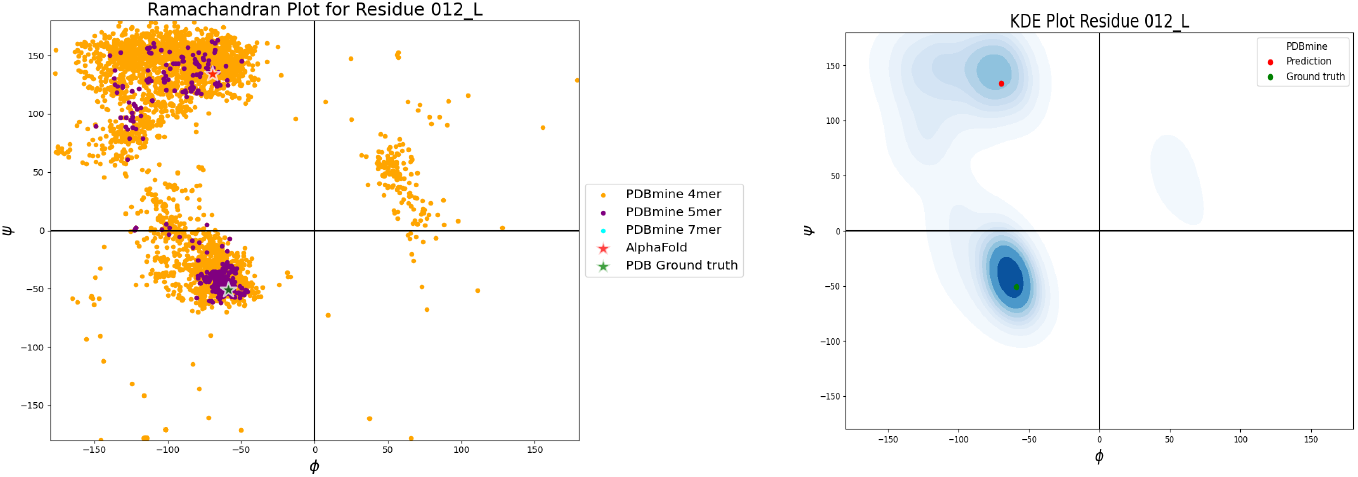
Ramachandran plots on the left depict PDBMine results for window sizes of 3mers (orange), 4mers (purple), and 7mers (cyan), with the X-ray torsion angle marked by a green star and the predicted AlphaFold2 model by a red star. The corresponding 3D projections on the right show the likelihood density, where PDBMine is represented as a density plot and experimental truth and prediction are marked by green and red points, respectively. 2a corresponds to T1032 residue 108 (isoleucine), and 2b to T1038 residue 12 (Glutamic acid).

**Fig. 3.**
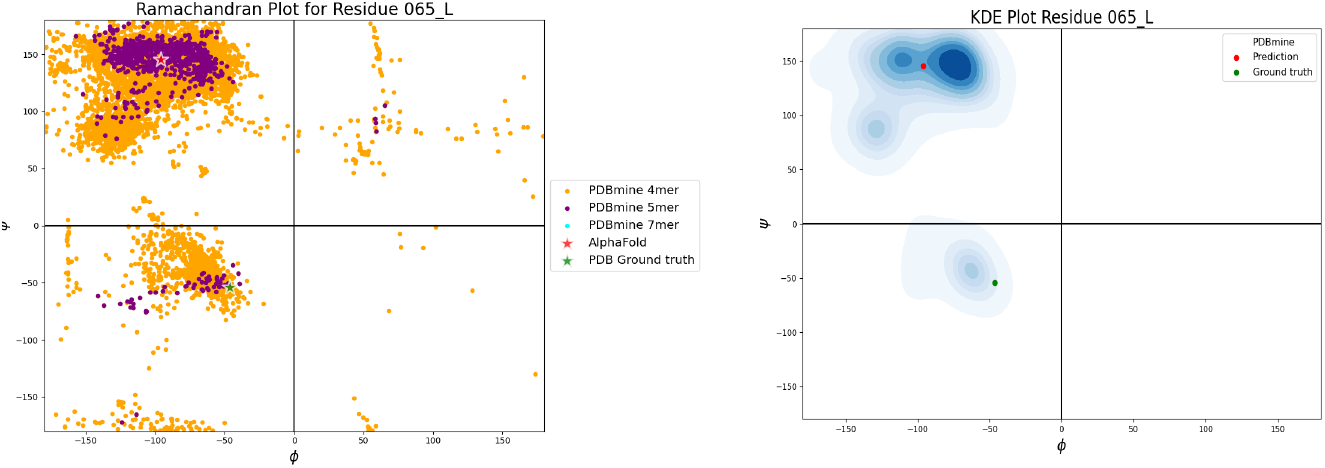
Ramachandran plots on the left depict PDBMine results for window sizes of 3mers (orange), 4mers (purple), and 7mers (cyan), with the X-ray torsion angle marked by a green star and the predicted AlphaFold2 model by a red star. The corresponding 3D projections on the right show the likelihood density, where PDBMine is represented as a density plot and experimental truth and prediction are marked by green and red points, respectively. Figure 3 corresponds to T1029-D1 residue 65 (lysine).

When PDBMine has limited data for a particular residue, the resulting sparse Ramachandran plot often leads to discrepancies, where the ground truth, PDBMine, and AlphaFold2 all disagree. This lack of data compromises the accuracy of the predictions. Additionally, certain unique cases arise where the ground truth coordinates deviate from the higher window size points generated by PDBMine. For example, in **Figure 3**, protein T1029-D1 residue 65 (lysine) shows that while the 5-mer cluster is densely concentrated in the beta-sheet region, the actual ground truth resides in a less populated alpha-helix domain cluster.

Likelihood analysis was conducted to determine which window size is more meaningful, as many residues exhibit multiple clusters. The 3D KDE (Kernel Density Estimation) was used to highlight these variations and provide a more precise depiction of the structural space. PDBMine yields valuable structural insights with potential protein structure prediction and refinement applications. By incorporating likelihood analysis, we can better assess each cluster, facilitating the selection of the optimal structural prediction for each residue.

### 4.2 Ideal peak vs. Considering other peaks

Using a binary mask based on 0.004% of the ideal peak from the 3D-KDE analysis, **Figure 4** depicts a stacked bar plot of the percent of acceptable regions for each residue of the target proteins under study. The lowest X-ray result for all the investigated proteins was used to determine the 0.004% criterion. Based on the data, each residue has an average of about 20% of its Ramachandran space categorized as acceptable. Specific residues demonstrate better flexibility and more extensive acceptable zones due to having more data points clustered in high-likelihood places. On the other hand, at least 20% of residues are accepted. It’s also crucial to notice that some proteins have white spaces, which represent the missing or gapped residues that are difficult to record because of their dynamic nature.

**Fig. 4a.**
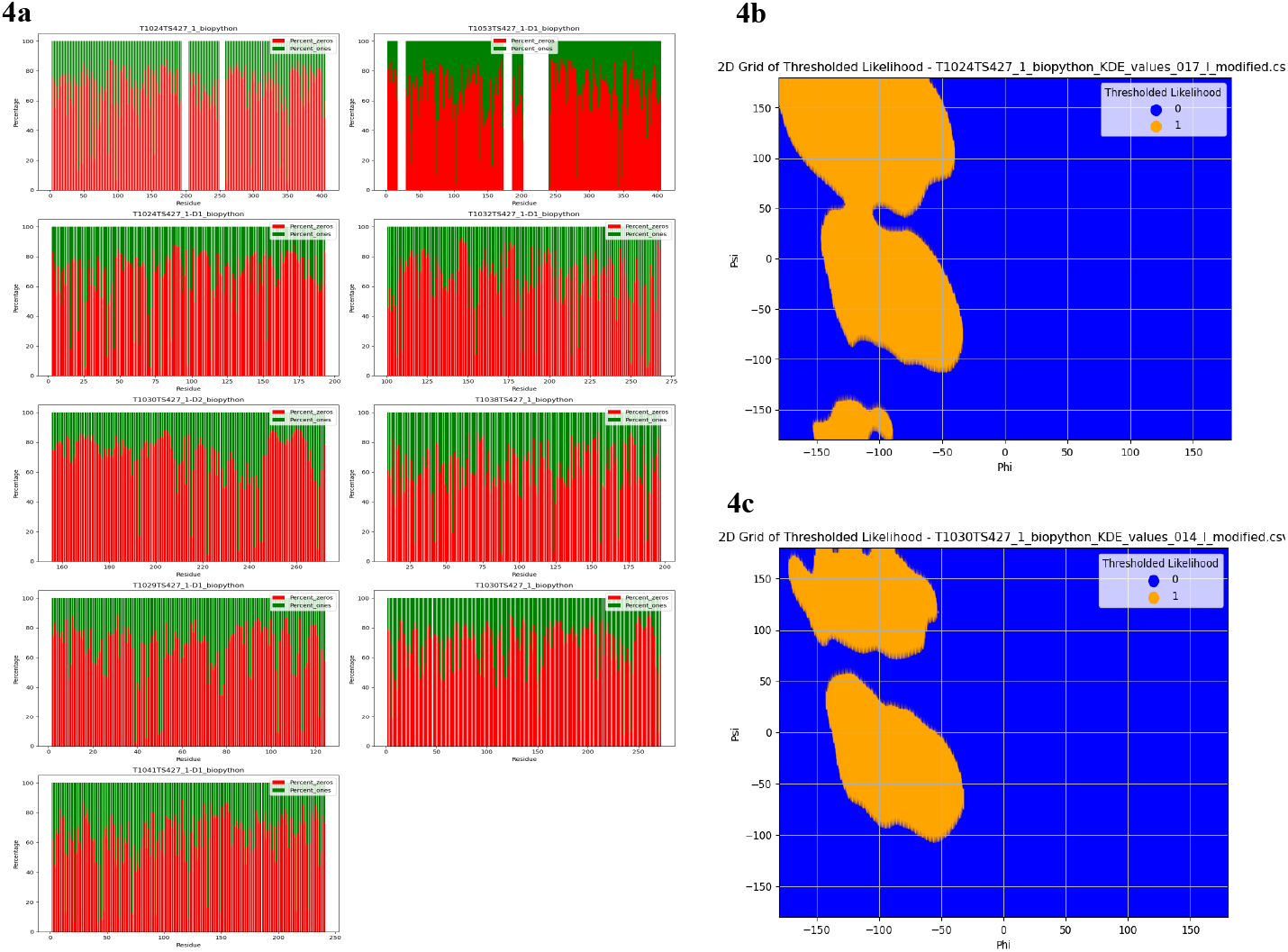
Stacked bar plot on the right, where the red bars represent the rejected regions, and the green bars represent the accepted regions of the binary mask. The plot visualizes the distribution of acceptance and rejection for each residue based on the 0.004% threshold derived from the 3D-KDE analysis. Fig. 4b and **Fig. 4c** illustrate the Ramachandran subspace, incorporating the 0.004% threshold to allow for additional acceptable regions. Both figures focus on Isoleucine residues; **Fig. 4b** represents T1024 residue 17, while **Fig. 4c** depicts T1030 residue 14. These visualizations highlight how the 0.004% threshold broadens these residues acceptable torsion angle regions, offering flexibility in structural prediction analysis.

The Ramachandran subspace has now been expanded to include additional potential clusters. In **Figure 4b** (T1024, residue 17), we observe that instead of focusing on a single cluster, the entire subspace now encompasses both the beta-sheet and right-handed alpha-helix regions of the Ramachandran plot, providing broader structural insights. Similarly, **Figure 4c** (T1030, residue 14) covers the beta-sheet region, offering a more detailed view of the structural conformation. This broader approach allows for a more flexible scoring metric. If a predicted torsion angle falls within the acceptable region, it is assigned a score of 1, while predictions outside the acceptable region receive a score of 0. For a protein with *n* residues, the closer the total score is to *n*, the more accurate the prediction. A perfect score of *n* would indicate an ideal prediction.

However, we encountered cases where the X-ray structure fell within regions with sparse data points, leading to specific residues being missed. To address this, we incorporated a neighboring point analysis to capture these residues better and enhance the robustness of the methodology. This extension of the approach provides a more comprehensive evaluation of structural predictions, even in less densely populated regions of the Ramachandran subspace.

### 4.3 Neighboring points Analysis

Euclidean distances were calculated between X-ray structures and PDBMine data, X-ray structures and predictions from CASP participants, and X-ray structures and AlphaFold2 predictions. **Figure 5a** presents a 1D-KDE distribution of the combined values for all target proteins, where most differences are concentrated below 50 degrees. However, some extreme deviations, ranging from 180 to 200 degrees, indicate residues that may have been incorrectly predicted. **Figure 5b** shows these profiles in subgraphs for individual target proteins, offering a more detailed view of the variations. Neighboring point analysis was applied to address these discrepancies, considering any predicted point within a 5-degree difference from the X-ray structure as valid. This adjustment improves the accuracy of predictions by capturing minor deviations that would otherwise be excluded.

**Fig. 5a.**
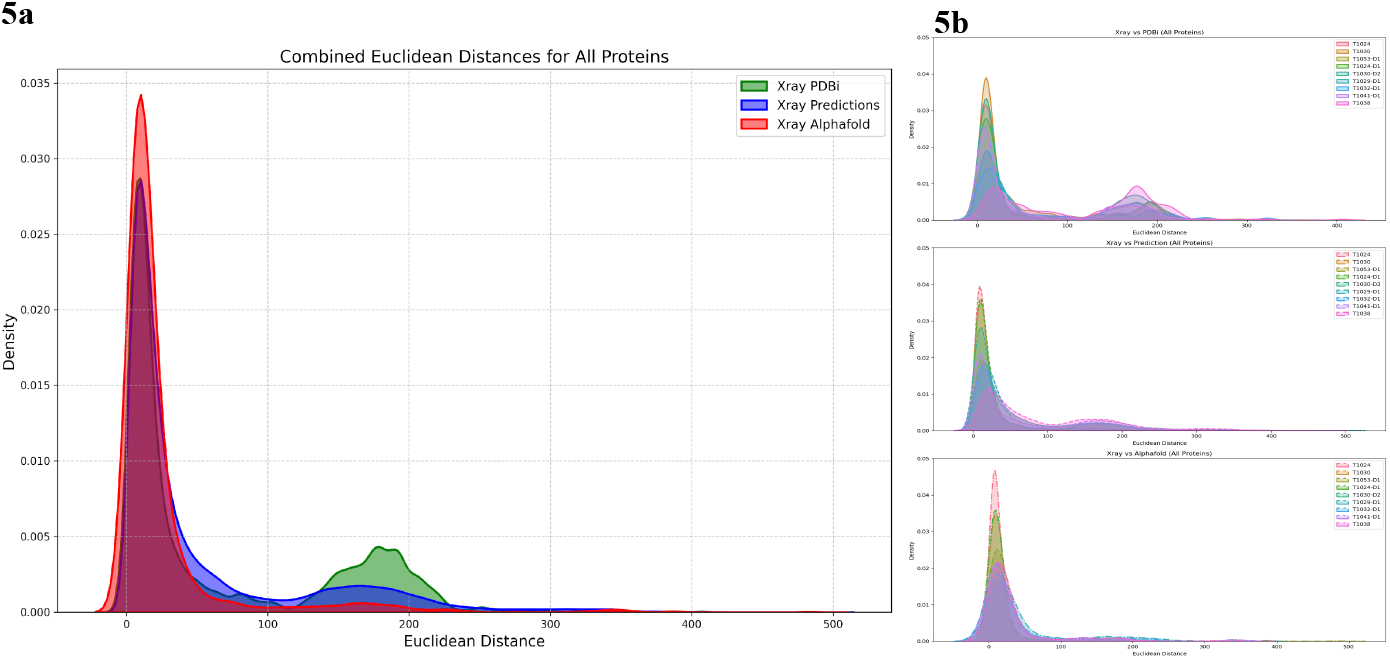
illustrates the 1D KDE distribution of Euclidean distances between the X-ray structures and predictions (CASP predictions in blue, AlphaFold2 in red, and PDBMine in green). The x-axis represents the Euclidean distance, while the y-axis shows the density. Fig. 5b presents similar subgraphs, separately highlighting the distribution of differences across all target proteins, comparing PDBMine predictions and AlphaFold2 results in sequence. These visualizations provide insights into the variation between predicted models and experimentally determined structures.

**Fig. 6.**
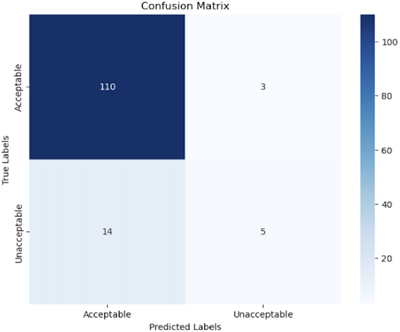
depicts the confusion matrix showing the performance of the decision tree classifier in identifying Acceptable and Unacceptable AlphaFold2 prediction models based on isolated points and RMSD on the Ramachandran plot. The model achieved an accuracy of 87% in detecting Acceptable models, with minor misclassifications observed primarily in identifying Unacceptable models.

This neighboring point analysis provided valuable insights. In 97% of cases, data points were found near the X-ray torsion angle, indicating strong concordance between predicted and experimental data. However, in a small but significant 3% of cases, the X-ray torsion angle did not correspond to previously observed experimental data points. These deviations were typically located in the protein’s gap regions or dynamic areas, where structural flexibility or variability is more pronounced. This analysis offered a deeper understanding of how torsion angles relate to sparsely populated areas within the Ramachandran subspace, particularly in structurally flexible regions.

The decision tree classifier, built to examine the structural dependability of AlphaFold2 prediction models based on isolated points within incremental radial thresholds on the Ramachandran plot, attained an overall accuracy of 87.17% using Leave-One-Out Cross-Validation (LOOCV). The classifier performed well in finding Acceptable models, with a precision of 0.89, a recall of 0.97, and an F1-score of 0.93, suggesting its ability to categorize reliable predictions. On the other hand, unacceptable models proved more challenging to classify, with a precision of 0.62, recall of 0.26, and an F1-score of 0.37, most likely due to the category’s lower support. According to the confusion matrix, the model correctly recognized 110 Acceptable models and 5 Unacceptable models but incorrectly classified 3 Acceptable models as Unacceptable and 14 Unacceptable models as acceptable. Despite these challenges, the classifier shows a strong capacity to discriminate structurally acceptable models based on the spatial distribution of isolated points, offering useful information for evaluating AlphaFold2 predictions.

## 5 Conclusion

To improve the precision of protein structure predictions, this study concludes that it is critical to incorporate various structural evaluation techniques, including nearby point analysis, 3D Kernel Density Estimation (KDE) profiles, and Ramachandran plots. We identified critical alignment regions and inconsistencies by comparing projected torsion angles from models like AlphaFold2 with experimental X-ray data and PDBMine. A more nuanced assessment of the predictions was made possible using variable scoring criteria based on phi and psi angles and a 0.004% threshold for cluster acceptance. This was especially useful when departures from the most likely torsion angle clusters occurred. More significant structural insights were obtained with larger k-mer window sizes, highlighting the importance of KDE profiles in capturing minute structural details.

The neighboring point analysis, which introduced a 5-degree tolerance around the X-ray torsion angles, played a pivotal role in addressing minor deviations, increasing the robustness of the predictions. This approach proved especially effective for residues in gap regions or sparsely populated areas, where traditional methods often need help. The methodology demonstrated the potential of flexible evaluation techniques in improving protein structure predictions and highlighted the need for advanced computational tools to refine future predictive models. Ultimately, these insights are crucial for driving progress in protein modeling and guiding therapeutic strategy development.

PDBMine significantly aids knowledge of protein torsion angles and structural preferences, which provides insightful structural information that can be directly applied to predicting and improving protein structures. This research improves computational modeling accuracy and expands our understanding of protein structure dynamics using clustering and likelihood analyses, computational methodologies, and experimental data. The capacity of PDBMine to extract and analyze structural data from the Protein Data Bank (PDB) is a potent tool for advancing our knowledge of protein geometry. It provides a robust framework for evaluating protein structure models computationally. In computational proteomics, this advancement improves the fidelity of structural models, facilitating more accurate protein structure predictions with enhanced biological significance.

